# Antiparasitic effect of peptoids against *Cryptosporidium parvum*

**DOI:** 10.1101/2024.07.16.603782

**Authors:** Bridget Lamont, Bruce M Russell, Rossarin Suwanarusk, Josefine Eilsø Nielsen, Kristian Sørensen, Deborah Yung, Annelise E. Barron, Daniel Pletzer, Morad-Remy Muhsin-Sharafaldine

## Abstract

Cryptosporidiosis, caused by *Cryptosporidium parvum*, poses significant health risks, particularly for children and immunocompromised individuals. Current treatments are ineffective in these vulnerable groups. This study explores the antiparasitic effects of against *C. parvum*. Out of 14 synthetic peptidomimetics (peptoids) screened, TM9 and TM19 exhibited potent anti-cryptosporidial activity without harming host cells. These findings suggest that peptoids could be a promising new therapeutic avenue for cryptosporidiosis, warranting further investigation.

---

Cryptosporidiosis, caused by the intestinal protozoan parasite *Cryptosporidium parvum*, causes diarrheal disease in humans and other mammals [1, 2]. This disease can result in life long symptoms and loss of life, particularly in children under five years of age or those who are immunocompromised [3, 4]. Despite its severe clinical manifestation for these groups, research into this parasite continues to be neglected and is made more difficult by the absence of an optimal *in vitro* continuous culture of the parasite. Furthermore, there are only two approved drugs to treat cryptosporidiosis in humans, nitazoxanide and paromomycin; however, neither of these drugs are effective in neonate infants or HIV-AIDs comorbidities [5, 6]. There is a pressing need for new and improved therapeutics for cryptosporidiosis. A common strategy for drug discovery involves investigating conserved drug targets identified in other Apicomplexan parasites, such as *Plasmodium* spp. However, inhibitors of these targets often encounter resistance, a prevalent and increasing issue with many therapeutics. Therefore, novel approaches are essential. [7]. Peptoids, synthetic mimetics of antimicrobial peptides, have shown potent inhibitory effects against a broad spectrum of bacterial and viral pathogens. Unlike traditional antimicrobials, peptoids are less prone to the rapid development of resistance, making them a promising avenue for novel therapeutic development [8, 9]. To date few studies have investigated the use of peptoids in parasites, all in *Leishmania* species [10-12]. In this study we investigate the antiparasitic effect of synthetic peptoids against *C. parvum*.

*Cryptosporidium parvum* oocysts obtained from the *Cryptosporidium* production laboratory (University of Arizona, USA) were kept in penicillin (100 U/mL) and streptomycin (100 µg/mL; ThermoFisher Scientific #15140122) in Phosphate-Buffered Saline (PBS, pH 7.2 (ThermoFisher Scientific #21600010)) at 4°C for no longer than four months. HCT-8, a human ileocecal adenocarcinoma cell line (ATCC # CCL-225) used as host cells, was gifted from the Guildford Lab (University of Otago, New Zealand). A library of 14 peptoids was synthesized by the Barron Lab (Stanford University, USA) and provided by the Pletzer Lab (University of Otago, New Zealand).

The library of 14 peptoids were diluted to five concentrations (5, 10, 20, 40 and 80 µg/mL) in RPMI 1640, no phenol red media (ThermoFisher Scientific #11835030) plus 3% horse serum (ThermoFisher Scientific #26050070). *C. parvum* sporozoites were released from the oocysts by pre-treating the oocysts with 1:4 dilution of household bleach for 10 minutes on ice, followed by incubating in 0.75% sodium taurocholate (Sapphire Bioscience #16214) for 45 minutes at 37°C. Parasites were added to confluent HCT-8 monolayers in 96-well plates at a concentration of 2 × 10^4^ oocysts per well and left to invade. After 3 hours, the media was removed and fresh media containing the peptoid was added and incubated for a further 45 hours. Each of the peptoid treated assay plates were harvested and stained at two timepoints; 3-hour (baseline control) and 48-hour (final efficacy). Plate wells were fixed with 4% paraformaldehyde (VWR Chemicals #28794.295) in PBS for 10 minutes at room temperature, permeabilised with 0.25% Triton X-100 (Merk #SLBP6453V) in PBS for 10 minutes at 37°C and blocked with 1% bovine serum albumin (Roche #3853143) in PBS for 1 hour at room temperature. The wells were then stained with 2 µg/mL Vicia Villosa-Fluorescein isothiocyanate (VVL-FITC; Vector Laboratories) in 1% BSA in PBS for 1 hour at room temperature in the dark, which stains the parasitophorous vacuoles (PV) of the parasite. This was followed by nuclei staining of the host cells with 2 µg/mL Hoechst 33342 in Milli-Q^®^ water for 15 minutes at room temperature in the dark. Between each stain the wells were washed with 0.1% Tween-20 (Merk #SLBZ8563) in PBS three times. The wells were then imaged on an Evos FL Auto 2 Imaging system microscope (Thermo Scientific Invitrogen EVOS) at 20 × magnification covering 35% of each well from the centre. The experiments were done in triplicate with three independent repeats and the doses were expanded for lead peptoids. The inhibitory concentrations were calculated using the ICEstimator version 1.2 (http://antimalarial-icestimator.net/).

Of the 14 peptoids screened, eight showed evident cytotoxicity to the host cell monolayer. TM9 and TM19 had the best antiparasitic effect, with minimal host toxicity and were selected for further investigation. We determined the IC50 and IC90 values as 21 µg/mL and 26 µg/mL, respectively, for TM9 and 27 µg/mL and 50 µg/mL, respectively, for TM19 (Figure 1).

**Figure 1:**
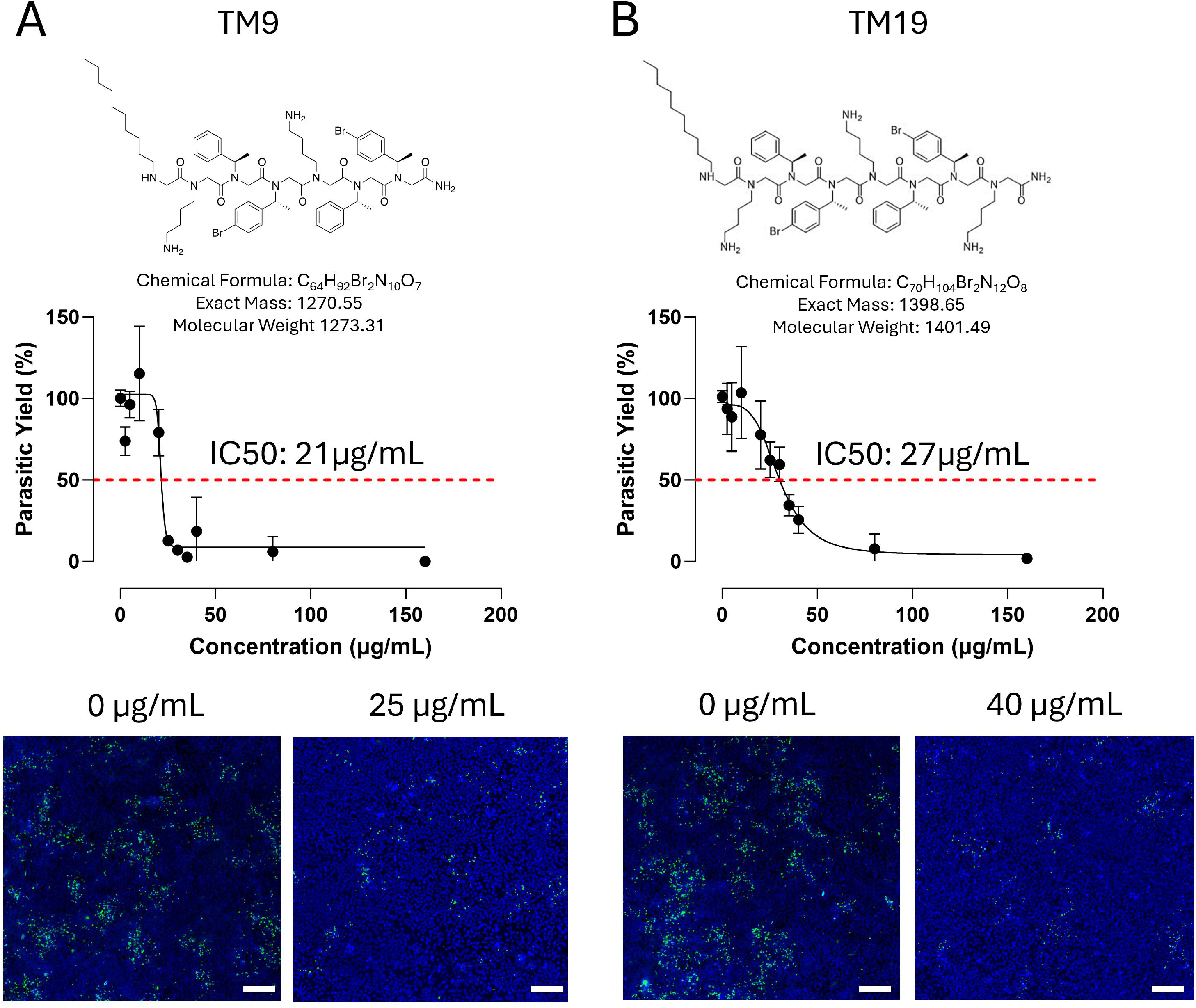
Structures and inhibitory concentration curves for lead peptoids targeting *Cryptosporidium parvum*. Chemical structures and Inhibitory concentration curves for TM9 (A) and TM19 (B). *C. parvum* infected HCT-8 cells were treated with concentrations of the peptoid ranging from 2.5 µg/mL to 160 µg/mL. At 48 hours post infection the wells were fluorescently stained with VVL-FITC and Hoechst 33342. The stained wells were then imaged using an Evos FL Auto 2 Imaging System Microscope and the parasitic yield was determined using Image J. The percentage parasitic yield was determined by normalising to the 0 µg/mL control well (PBS). Results are pooled from three independent experiments. Representative images of *C. parvum* infected HCT-8 cells not treated with peptoid (0 µg/mL) and treated with 25 µg/mL of TM9 or 40 µg/mL of TM19. Photos taken at 20 X magnification. Scale bar = 100 µm.

The two lead peptoids TM9 and TM19 are structurally almost identical, with TM19 containing one additional lysine-like residue. TM9 and TM19 self-assemble, predominantly into ellipsoidal core-shell micelles, which has been found to play an important role in their antimicrobial activity in a study on *ESKAPE* pathogens [13]. Bromination of these peptoids may have enhanced their anti-parasitic activity, as halogenation, particularly with chlorine or bromine, has been shown to increase the antimicrobial effects of peptoids [14]. The mechanism of action for peptoids against bacteria such as *Escherichia coli*, is suggested to be damage to the bacterial cell membrane [15] or membrane permeabilization followed by intracellular flocculation of biomacromolecules including ribosomes [16]. A similar mechanism is inferred for viruses such as Herpes Simplex Virus 1 and SARS-CoV-2 where peptoids, including TM9(MXB-9), affected the viral envelope [9]. Additionally, TM9 has been shown to disrupt the viral envelopes of Zika and chikungunya viruses [17] while the reported anti-hepatitis B virus effects of TM19 are also likely due to disruption of the viral envelope [18]. In the case of Leishmania, the only other parasite examined for its susceptibility to peptoids, it was shown that these compounds (notably TM9) preferentially target the promastigote membrane, which is composed of polysaccharide lipophosphoglycan, while not affecting the host cell membrane that carries phosphatidylcholine and sphingomyelin on its outer surface [12]. It is, therefore, reasonable to suggest that a similar mechanism may be occurring with Cryptosporidium parvum, wherein the peptoids might damage the parasitophorous vacuole. However, electron microscopy studies are warranted to validate this theory. Given the complexity of the parasite’s life cycle, it is also essential to test peptoids on different life cycle stages, including invasion, merozoite egress, and gamete formation. Nonetheless, we have demonstrated, for the first time, that peptoids can exhibit anti-cryptosporidial activity and, with some modifications, present a promising option for new and improved treatment for cryptosporidiosis

## Acknowledgements

BL, BR, RS, and RM were supported by a 2021 University of Otago Research Grant and a 2021 MBIE Science Whitinga Fellowship.

